# An efficient, scarless, selection-free technology for phage engineering

**DOI:** 10.1101/2023.03.26.534254

**Authors:** Moran G Goren, Tridib Mahata, Udi Qimron

## Abstract

Many phage engineering technologies, mainly based on the CRISPR-Cas system, have been recently developed. Here we present a method to genetically engineer the *Escherichia coli* phages T5, T7, and P1 by adapting a technology, called pORTMAGE, developed for engineering bacterial genomes. The technology comprises of *E. coli* harboring a plasmid encoding a potent recombinase and a gene transiently silencing a repair system. Oligonucleotides with the desired phage mutation are electroporated into *E. coli* followed by infection of the target bacteriophage. The high efficiency of this technology, ranging from 1-14% of desired recombinants, allows low-throughput screening for the desired mutant. We demonstrated the use of this technology for single-base substitutions, for deletions of 50 bases, for insertions of 20 bases, and for three different phages. The technology may be adjusted for use across many bacterial and phage strains.

## Introduction

The abundance of phages and their importance to microbial evolution and major ecological issues provide an incentive to study their biology^1^. In recent years, several methods have been developed to engineer phage genomes using retroelements^2^ and heterologous supplement of recombination enzymes^3^. We were the first group to harness the CRISPR-Cas system for the genetic engineering of phages^4^, followed by many others^5-10^. Using CRISPR-Cas for genetic engineering is straightforward, can be designed for virtually any gene, and is cost-effective. However, this technology requires specific plasmids for each genetic manipulation and is restricted to host cells encoding a functional CRISPR-Cas system.

Recently, new technologies have emerged, showing efficient engineering of many genomes, including bacterial genomes, but these were not adapted for phage engineering^11,12^. One of these technologies, the pORTMAGE (Parallel Oligonucleotide Recombineering and Targeted Multiplex Genome Editing), enables the simultaneous and efficient modification of multiple sites in a genome by introducing mutated oligonucleotide along with the inactivation of a critical repair protein, leading to homology-directed repair. Bacterial mutagenesis using pORTMAGE has been successfully shown, with lambda red^11^ and later with the significantly improved recombination enzyme CsRecT^12^. Bacteriophage engineering may use these similar principles but requires adaptation of each step, such as protein induction, electrocompetent preparation, growth media, and the timing and the multiplicity of infection.

Here we describe a simple and efficient method to genetically engineer the *E. coli* phages T5, T7, and P1. We term it pORTPHAGE, and use it to generate the desired mutations at high efficiencies. The identification of the recombinants is carried out by simple Sanger DNA sequencing or by PCR. We show by random sampling of the recombinants and by next-generation sequencing (NGS) that the efficiency of the technology enables low-throughput screening for the isolation of desired mutants from all tested phages. We show that the technology allows the generation of point mutations, insertions, and deletions. This method could be easily adjusted to other bacterial hosts, with minor modifications, as the pORTPHAGE plasmid is universally designed to function in many bacterial strains.

## Results

### The pORTPHAGE system

To develop pORTPHAGE (a revised pORTMAGE method^11^ adapted for phages), an efficient selection-free and scarless technology for genetically engineering various phages, we used *E. coli* as the host in which the different genetic manipulations were carried out. Into this host, we introduced the pORTPHAGE plasmid - pORTMAGE-Ec1^12^. This plasmid encodes the recombination enzyme RecT, with the highest reported recombination efficiency, originating from a phage infecting *Collinsella stercoris*. It also encodes a dominant-negative allele of the MutL protein, which temporarily silences the repair protein MutL and thus prevents the repair of the introduced mutation^13^. We speculated that bacteria expressing the above proteins from the plasmids, infected with phages, and electroporated with oligonucleotides encoding mutations, will produce desired recombinant phages. We also speculated that the high efficiency of this procedure would allow affordable screening for identifying the desired recombinants (Figure 1).

**Figure 1.**
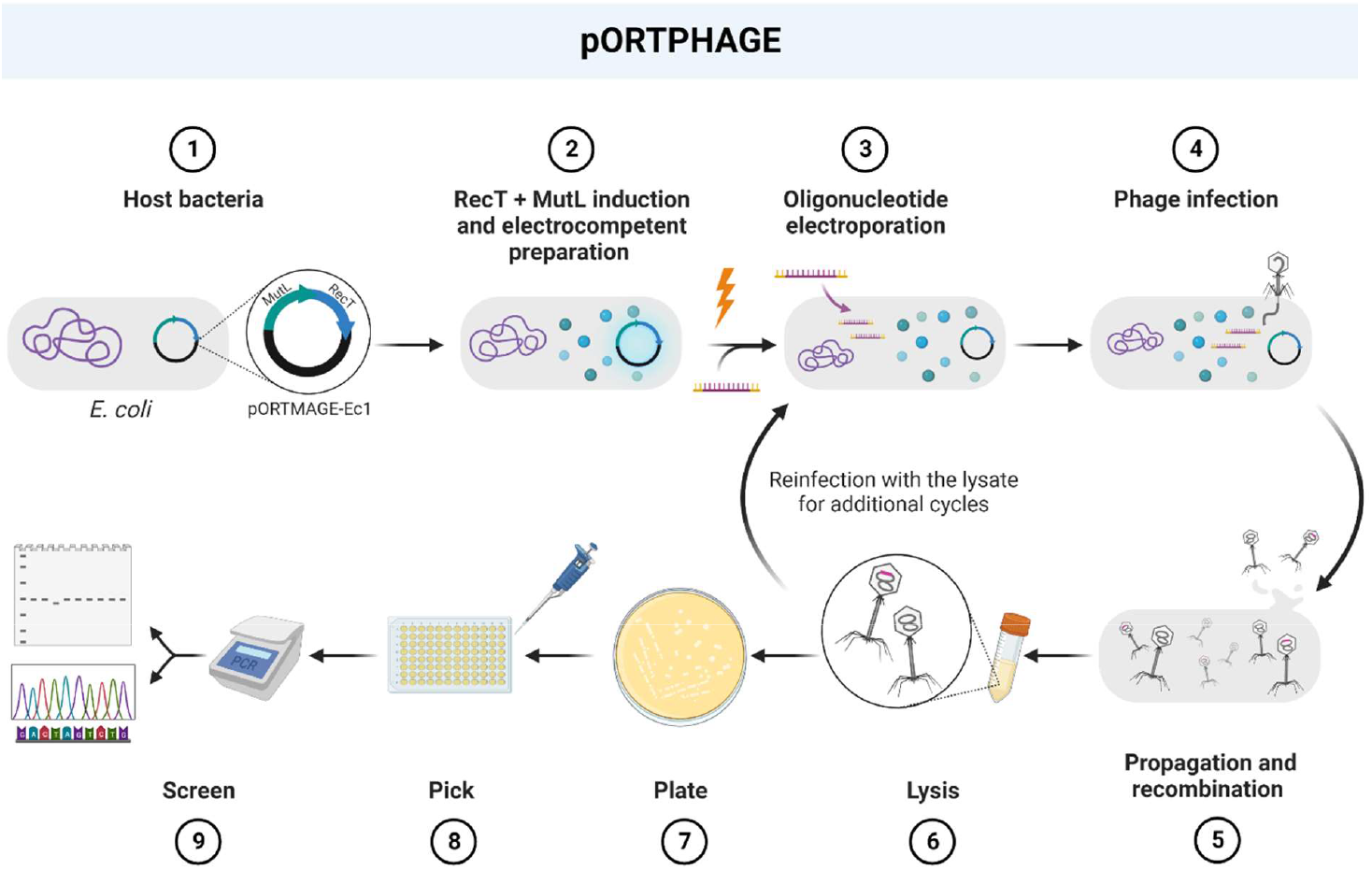
Schematic representation of the pORTPHAGE steps. **1**. Transformation of pORTMAGE-Ec1 into the host bacteria. **2**. Bacterial growth to early log phase followed by expression of recombineering proteins and preparation of electrocompetent bacteria. **3**. Electroporation of oligonucleotides with desired mutation. **4**. Infection of bacteria with the bacteriophage target. **5**. Phage propagation and recombination. **6**. Obtained lysate used for an additional cycle (step 3) or: **7**. Obtained lysate grown on bacterial host. **8**. Single plaques are individually picked. **9**. Relevant segments amplified by PCR and analyzed on agarose gel or Sanger sequencing.

### Generation of point mutations in T7 phage

We first tested whether we could generate a point mutation in a T7 bacteriophage. We chose a T7 phage with an amber mutation in gene 1, encoding T7-RNA polymerase, an essential protein for T7 phage propagation. This mutant phage can grow on amber-suppressor host, such as NEB5α, but not on the wild-type K-12 *E. coli*. This phenotype allows an easily detectible assessment of the recombination efficiency. We separately electroporated two oligonucleotides encoding a correction of the amber mutation in two reverse-complement orientations (which we arbitrarily term F and R) into an amber-suppressor host harboring pORTMAGE-Ec1, and then infected it with the amber-mutated T7 phage. We carried out a similar procedure as a negative control but used water instead of oligonucleotides for electroporation. Following two cycles, illustrated in Figure 1, we plated serial dilutions of the lysates, and replica-plated 96 plaques from each lysate on the K-12 and the NEB5α amber-suppressor host. We then measured the number of corrected phages for the treated versus untreated phage on K-12 *E. coli* compared to the amber suppressor host. In this way, we detected 11/96 of corrected phages for one oligonucleotide, 4/96 for the complementary nucleotide, and 3/96 of spontaneous revertants in the untreated control sample. This result suggests that the pORTPHAGE mutated on average ∼5% of the phage progeny after subtracting the background. Most treated recombinants (9/13) that grew on the wild-type host showed the expected correction in Sanger sequencing. The procedure thus enables the isolation of desired mutations at high frequencies.

### Validation by NGS and by Sanger sequencing

To evaluate the mutagenesis efficiency by an unbiased and statistically robust methodology, we carried out NGS on PCR-amplified segments from gene 1, encoding the mutation, which were grown on the amber-suppressor host without positive selection for the mutation. Analysis of the NGS showed that the mutation occurred in 4.5% for one oligonucleotide (F) and 7.9% for the other oligonucleotide (R), corroborating the results obtained above.

### Point mutations in different phages

We used the same protocol for mutating another gene of T7 phage, gene 17. In this case, we identified the desired mutant in ∼3% using Sanger sequencing for one oligonucleotide (R), and 0% for another. The NGS showed higher efficiency of ∼7% and ∼2%, respectively (Table 1). This suggests that the orientation of the oligonucleotides affects the outcome^14^, and both should be used for the procedure.

**Table 1.** Point mutations obtained using pORTPHAGE in T7 and T5 phages.

| Phage | Host harboring<br>*pORTMAGE-Ec1* | Targeted gene | Oligonucleotide<br>orientation | Recombinants<br>identified by<br>Sanger<br>sequencing (%) | Recombinants<br>identified by NGS<br>(%) |
| --- | --- | --- | --- | --- | --- |
| T7 | NEB5α | 1 | F | 2.08 | 4.49 |
|  |  |  | R | 7.29 | 7.87 |
|  |  | 17 | F | 0 | 2.01 |
|  |  |  | R | 3.13 | 6.90 |
|  |  | control | no | ND | 0 |
| T5 | DH5α | D15 | F | 0.5 | 0.9 |
|  |  |  | R | ND | 1.02 |
|  |  | control | no | ND | 0.05 |

We next used a slightly modified protocol for mutating a T5 phage. We applied 4 cycles before screening (Figure 1) due to the inconsistent efficiency of the mutagenesis of this phage^2^. The Sanger screening yielded 0.5% of recombinants only for one of the oligonucleotides (F) and 0% for the other (R), whereas the NGS showed 0.9% and ∼1% recombinants, respectively. The controls, with no oligonucleotides, yielded negligible percentages of mutations in the NGS. These results demonstrate that we can use the protocols for easily screening the desired recombinants without the requirement to establish a negative selection approach such as CRISPR-Cas.

### Deletions in different phages

Encouraged from these results, we carried out similar protocols on T7 and P1 for deleting segments of 50 bp. We used PCR to easily detect the band shifts stemming from deletions and insertions. We found that for T7, the PCR of individually isolated plaques showed 5-6.25% recombinants whereas for P1 phage, 2-4.9% (Table 2). The NGS in all cases showed even higher frequencies of mutagenesis. These results demonstrate that deletions can be obtained with high efficiency and easily screened using PCR.

**Table 2.**
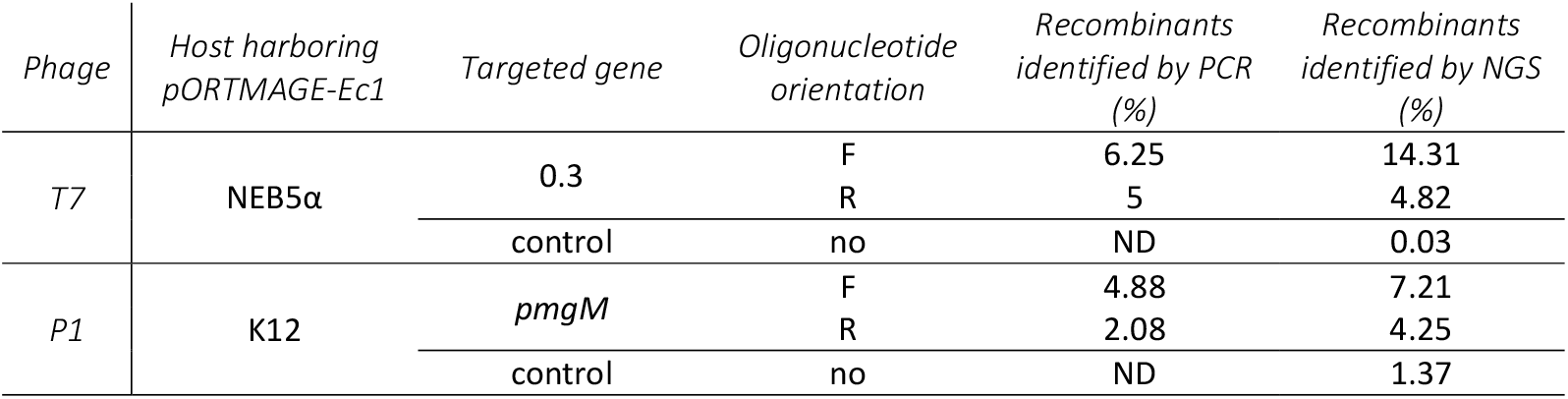
Deletion obtained by pORTPHAGE in T7 and P1 phages.

### Insertion of 20 bp in T7 phage

Finally, we wished to demonstrate that the procedure is also effective for insertions. We used a similar protocol to insert 20 bp segments into T7 phage, with oligonucleotides encoding the insertion. Here too, we used four cycles, obtaining for one oligonucleotide ∼4% recombinants with the desired insertion as detected by PCR, and none for the other oligonucleotide. NGS showed a 2% frequency for the first oligonucleotide and 1% for the other (Table 3). These results demonstrate that insertions can be obtained with high efficiency and easily screened using PCR.

**Table 3.** An insertion obtained by pORTPHAGE in T7 phage.

| *Phage* | *Host harboring*<br>*pORTMAGE-Ec1* | *Targeted gene* | *Oligonucleotide*<br>*orientation* | *Recombinants*<br>*identified by PCR*<br>*(%)* | *Recombinants*<br>*identified by NGS*<br>*(%)* |
| --- | --- | --- | --- | --- | --- |
| T7 | NEB5α | 0.3 | F | 4.17 | 2.21 |
|  |  |  | R | 0 | 1.02 |
|  |  | control | no | ND | 0 |

## Discussion

The technology shown here for the genetic engineering of phage genomes is adapted from a recent technology for bacterial genome engineering. Since the phage cannot harbor the plasmid, and relies on a bacterial host for propagation and recombination, it is required to optimize the timing of the infection with the induction of the recombination\repair silencing proteins and the electroporation of the oligonucleotides. We demonstrate here a system that synchronizes these events and provides robust mutagenesis efficiencies. A universal genetic engineering system to engineer phages may prove highly valuable for phage research and biotechnological use.

We used the most efficient recombination enzyme described to date, resulting in mutagenesis frequencies of between 1% and 14%. These frequencies allow for easy screening of the desired mutants. The technology is easy to transfer and maintain in the lab – requiring only a bacterial host harboring one plasmid, and using commercially available 5’ and 3’-modified oligonucleotides. The mutagenesis procedure is straightforward, allowing the generation of several genetically engineered loci in less than a week of work. Indeed, we use the technology developed here routinely for engineering phages in our lab (e.g., Mahata et al, in preparation). We believe that with few adjustments, the system could easily be adapted for manipulating any phage genome in any host strain.

### Materials and methods

### Reagents, strains, and plasmids

Lysogeny broth (LB) medium (1% w/v tryptone, 0.5% w/v yeast extract, 0.5% w/v NaCl) was from Condalab, 2YT (1.6% w/v tryptone, 1% w/v yeast extract, 0.5% w/v NaCl) was from Neogen, Terrific broth (TB) was from Sigma-Aldrich, Agar was from Hylabs. MgSO_4_, CaCl_2_, Kanamycin and M-Toluic acid was from Merck. KAPA HiFi HotStart Ready Mix was from Kapa Biosystems. PCRBIO HS Taq DNA polymerase was from PCR Biosystems. NucleoSpin Gel and PCR Clean-Up Kit were from Geneaid. All PCR Standard oligodeoxynucleotides (oligos) were ordered with standard desalting from Merck. pORTPHAGE recombineering oligos containing two phosphorothioate (PT) bonds, in each end (3’ and 5’, marked by asterisks) were ordered with standard desalting from Integrated DNA Technologies (IDT). Bacterial strains, plasmids and oligonucleotides used in this study are listed in Supplementary Table S1.

### Detailed pORTPHAGE protocol

A single colony from a strain harboring the pORTMAGE-Ec1 plasmid was inoculated in LB medium supplemented with 50 μg/mL kanamycin and aerated in a shaking incubator at 37°C, 250 rpm for 16 hours (h). Overnight (ON) cultures were diluted 1:100 into LB medium containing 50 µg/mL kanamycin and incubated at 37 °C, 250 rpm. Upon reaching OD_600_ ∼0.3, cells were induced for RecT expression with 1 mM m-toluic acid for 30 min at 250 rpm. After induction, cells were immediately chilled on ice for at least 10 min and made electrocompetent by washing and pelleting twice in 10 mL of ice-cold dH2O at 4,800 × g for 10 min in a chilled Thermo X4R Pro centrifuge with swing-out rotor. Electrocompetent cells were suspended in 160 μL of ddH2O and were kept on ice until electroporation. 45-μL cell suspension was mixed with 5 μL of 100 μM oligonucleotide and Electroporated using BioRad MicroPulser in pre-chilled 2-mm gap Electroporation cuvettes (1.8 kV, 200 Ω, 25 μF). Immediately after electroporation, 1 mL of room-temperature (RT) phage infection media was added and the suspension was then transferred to 4 mL RT media supplemented with phage lysate at an MOI of 1:1. Each phage was supplemented with different infection media. For T7 we used TB, for T5 we used LB supplemented with 10 mM MgSO_4_ + 10 mM CaCl_2_, and for P1_vir_ we used LB supplemented with 5 mM CaCl_2_. Cells and phage mixture were incubated at 37°C under continuous agitation until culture lysed completely. 100 µL of chloroform were then added to the lysates, mixed and centrifuged at 4,800 × g for 10 min. The lysates were used for further rounds of infections as indicated. When the last cycle reached its end-point, it was used as a PCR amplification template for NGS, then serially diluted and plated to screen for individual recombineered plaques. For NGS, 1ul from the lysate was used as template for 20 cycles of PCR amplification using KAPA HiFi polymerase and primers containing different 8 nucleotide tags (marked as X) for each sample. Amplified products were purified using PCR cleanup kit and sent for NGS. For single plaque screens, the lysate was serially diluted (10^−1^-10^−7^) and plated on a permitting host using the soft-agar overlay technique. The plates were incubated ON at 37°C for T5, and P1 or at RT for T7. Single plaques were picked, suspended in 20 µL of LB and used as template for PCR amplification using Taq polymerase and the same primers used for the NGS, but without the tags. Amplified products were purified and sequenced using the Sanger method with one of the primers used for amplification.

### NGS Library preparation

The PCR products were purified and used as templates for a second PCR to barcode the samples specifically and enable NGS readings. The prepared Illumina sequencing libraries were sequenced in Novogene using NovaSeq 6000 machine according to the manufacturer’s instructions, using 2×150 flow cell. Samples were multiplexed in the same sequencing run. Demultiplexing was based on a 8-bp barcode that was part of the original PCR primer.

### NGS data analysis

The input is composed of paired-end sequencing. Each end is 150 bases long. The import program looks for an overlap between the end of the forward read and the beginning of the reverse complement of the backward read and generates the combined sequence. Each generated sequence is grouped according to one of its given primers. Sequences are then divided into subgroups of equal length. For each subgroup and each location in the sequence, each nucleotide (A, C, G, T) is counted (=histogram). Gaps will be counted as “Deleted”. In the other locations, histograms are summed.

## Acknowledgements

UQ is supported by the European Research Council – Horizon 2020 research and innovation program, grant no. 818878. UQ has also received funding from the Israeli Ministry of Health in the framework of the ERANET-JPI-AMR (grant no. 15370).

## Supplementary Material

**Table S1.**
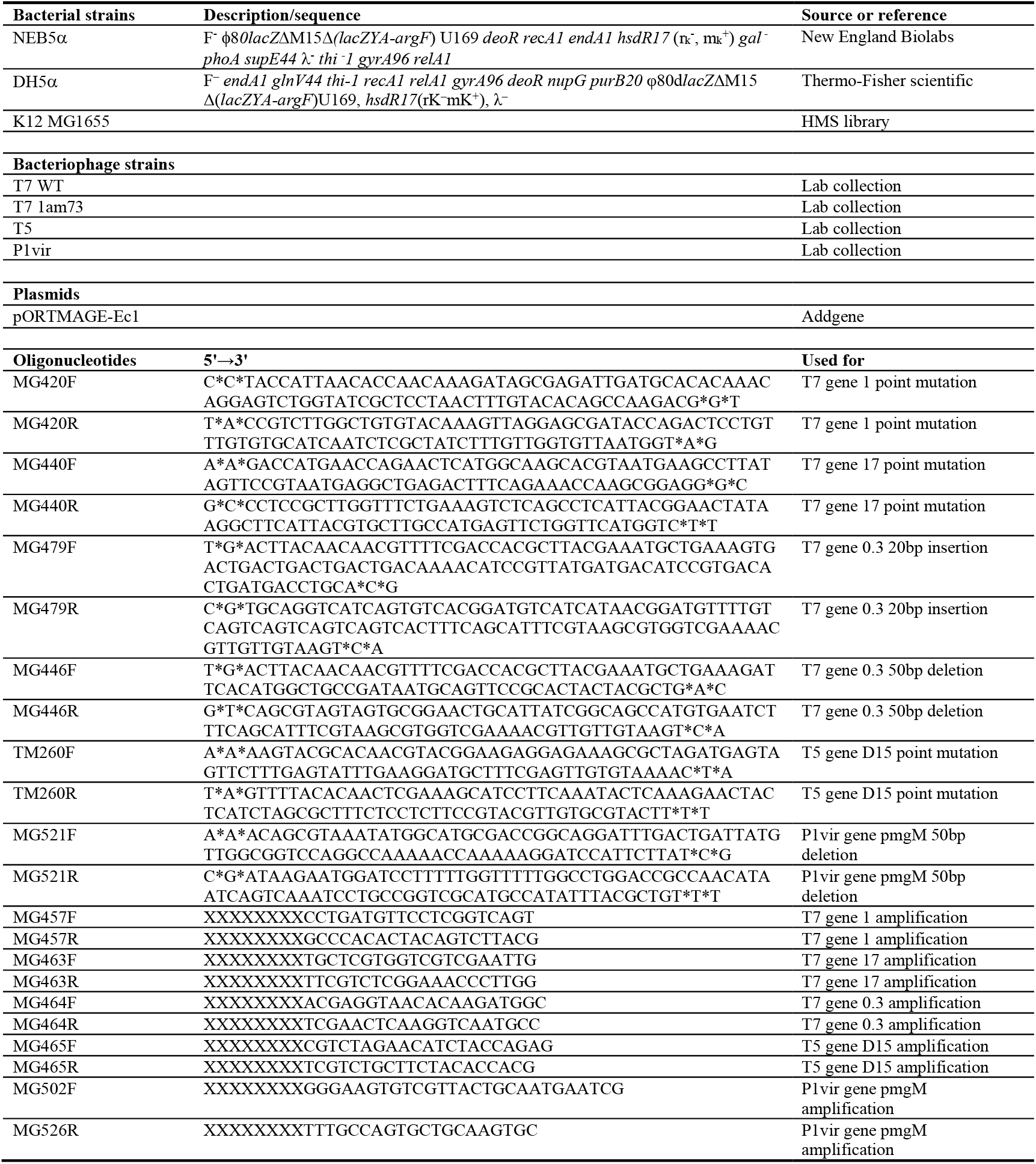
Bacterial strains, plasmids and oligonucleotides used in this study. Related to Experimental Procedures.

